# Transcriptome analysis reveals the genetic basis underlying the development of skin appendages and immunity in hedgehog (*Atelerix albiventris*)

**DOI:** 10.1101/2020.05.09.086496

**Authors:** Hui-Ming Li, Bi-Ze Yang, Xiu-Juan Zhang, Hai-Ying Jiang, Lin-Miao Li, Hafiz Ishfaq Ahmad, Jin-Ping Chen

**Affiliations:** Guangdong Key Laboratory of Animal Conservation and Resource Utilization, Guangdong Public Laboratory of Wild Animal Conservation and Utilization, Guangdong Institute of Applied Biological Resources, Guangdong Academy of Science, Guangzhou 510260, China

**Keywords:** hedgehog, skin appendage, adaptive evolution, RNA-Seq, molecular biology

## Abstract

The expression of hair features is an evolutionary adaptation resulting from interactions between many organisms and their environment. Elucidation of the mechanisms that underlie the expression of such traits is a topic in evolutionary biology research. Therefore, we assessed the *de novo* transcriptome of *Atelerix albiventris* at three developmental stages and compared gene expression profiles between abdomen hair and dorsal spine tissues. We identified 328,576 unigenes in our transcriptome, among which 3,598 were differentially expressed between hair- and spine-type tissues. Dorsal and abdomen skin tissues 5 days after birth were compared and the resulting differentially expressed genes were mainly enriched in keratin filament, epithelium cell differentiation, and epidermis development based on GO enrichment analysis, and tight junction, p53, and cell cycle signaling pathways based on KEGG enrichment analysis. Expression variations of *MBP8, SFN, Wnt10, KRT1,* and *KRT2* may be the main factors regulating hair and spine differentiation for the hedgehog. Strikingly, DEGs in hair-type tissues were also significantly enriched in immune-related terms and pathways with hair-type tissues exhibiting more upregulated immune genes than spine-type tissues. Thus, we propose that spine development was an adaptation that provided protection against injuries or stress and reduced hedgehog vulnerability to infection.

## Introduction

Perceiving and responding to life-threatening signals and regulating their own morphological characteristics constitute a fundamental challenge for all mammals. Evolution has shaped the ability of organisms to adapt skin appendages to increase their chances of survival. Accordingly, a variety of appendages have evolved on the skin of mammals, either to assist organisms in carrying out specific behaviors or to protect them from predators and pathogenic bacteria^1,2^.

Compared with animals that possess passive defense adaptations such as feathers, scales, etc., the hedgehog has evolved into an unusual, nocturnal, spine-covered mammal^3^. Spines potentiate attacking power and enhance the role of skin appendages as a defense mechanism. The hedgehog progresses through a series of well-defined stages during its life cycle, from embryonic, to non-spine, to with-spine stages. These transitions are governed by tightly regulated gene expression at pre-transcriptional, epigenetic, and translational levels. Spine- and hair-type skin tissues from the hedgehog thus offer a natural model to analyze the genomic basis for the evolution of epidermal appendage formation.

Development of hair follicles on a skin appendage with a complex growth cycle is characterized by anagen, catagen, and telogen stages, and many key signaling pathways and genes are involved in their regulation^4–6^. According to many studies, the WNT/b-catenin signaling pathway in the dermis may be the first dermal signal^7–9^. Sonic hedgehog (SHH) is another secreted protein in the follicular placode that plays a major part in epithelial-mesenchymal signaling^10,11^. SHH depends on WNT signaling and is required for the proliferation of follicular epithelium and development of dermal condensate into dermal papilla^4^. In addition, many key genes related to skin appendage development have been identified; *TGFaR* (epidermal growth factor receptor) and transcription factor *ETS2* regulate hair follicle shape and are responsible for hair follicle architecture and wavy hair^12,13^, and *HOXC13* and *KRT75* control hair shaft differentiation^14–17^. Further, *EDA* and *EDAR* interact with members of the bone morphogenetic protein (BMP) family, some of which inhibit follicle development to establish follicle patterning^2,18–20^.

The abovementioned studies primarily focused on mammalian hairs and scales; little is known about changes in gene expression during the development of spine appendages in mammals. Therefore, we aimed to perform a detailed analysis of the transcriptome of skin from abdomen hair and dorsal spines of the hedgehog (*Atelerix albiventris*) by comparing groups of transcripts differentially expressed at different hedgehog developmental stages. We identified key candidate genes related to the development and differentiation of skin appendages whose importance should be verified in future studies. Nonetheless, the presented novel information will be widely applicable in many fields, such as skin disease and skin immune genes, and provide insight into the molecular mechanisms of skin appendage development in general.

## Results

### Illumina sequencing and *de novo* assembly

To identify the transcriptome and molecular mechanisms governing skin appendage development and differentiation, we analyzed temporal changes in transcript abundance of *A. albiventris* (Fig. 1a). RNA sequencing generated 82.2 ± 6.0 G (mean ± SD) read pairs for 16 skin tissue samples (Supplementary Table 1). After quality trimming, 96.8 ± 0.4% of reads were retained, indicating a high-quality dataset (>90% reads with ≥Q30). The *de novo* assembly using Trinity revealed 328,576 unigenes, which were used for subsequent analysis. We evaluated biological reproducibility by individually comparing biological replicates through principle component analysis (PCA) (Fig. 1b). As expected, similar gene expression patterns in both hair- and spine-type samples were observed. PCA analysis also demonstrated more separation between the two skin appendage types. A neural network graph based on self-organizing feature map (SOM) analysis revealed dynamic transcriptional changes at the individual stages of appendage development in *A. albiventris* (Fig. 1c).

**Figure 1.**
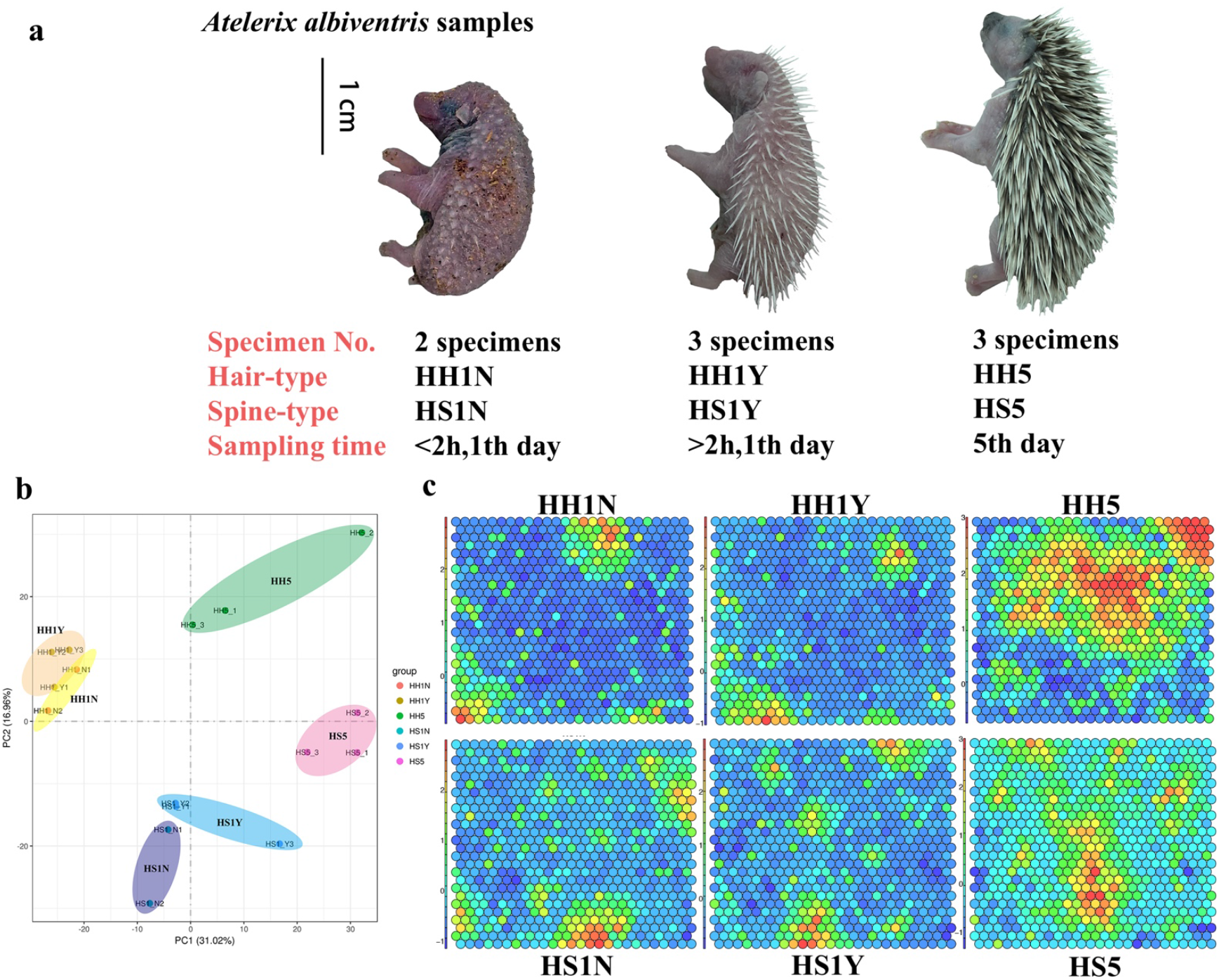
Phenotypes of skin appendages from 3 developmental stages of *Atelerix albiventris*. (a) Different phenotypes from 3 different developmental stages of *A. albiventris*; (b) Principal components analysis (PCA); (c) Gene expression-specific and phenotype-specific gene-trait correlation analysis based on self-organizing feature map module analysis.

### Functional annotation and classification of unigenes

Use of the NR and Swiss-Prot databases yielded reliable protein annotations for approximately 42% of unigenes. Absolute unigenes were annotated using seven major public protein databases (Table 1), of which the greatest number of matches (68.3%) was obtained using the NT database. Of 328,576 unigenes assembled, 243,444 genes (74.1%) exhibited a positive match against at least one database. Thus, our gene annotation results from *A. albiventris* transcriptome were considered high quality.

**Table 1.**
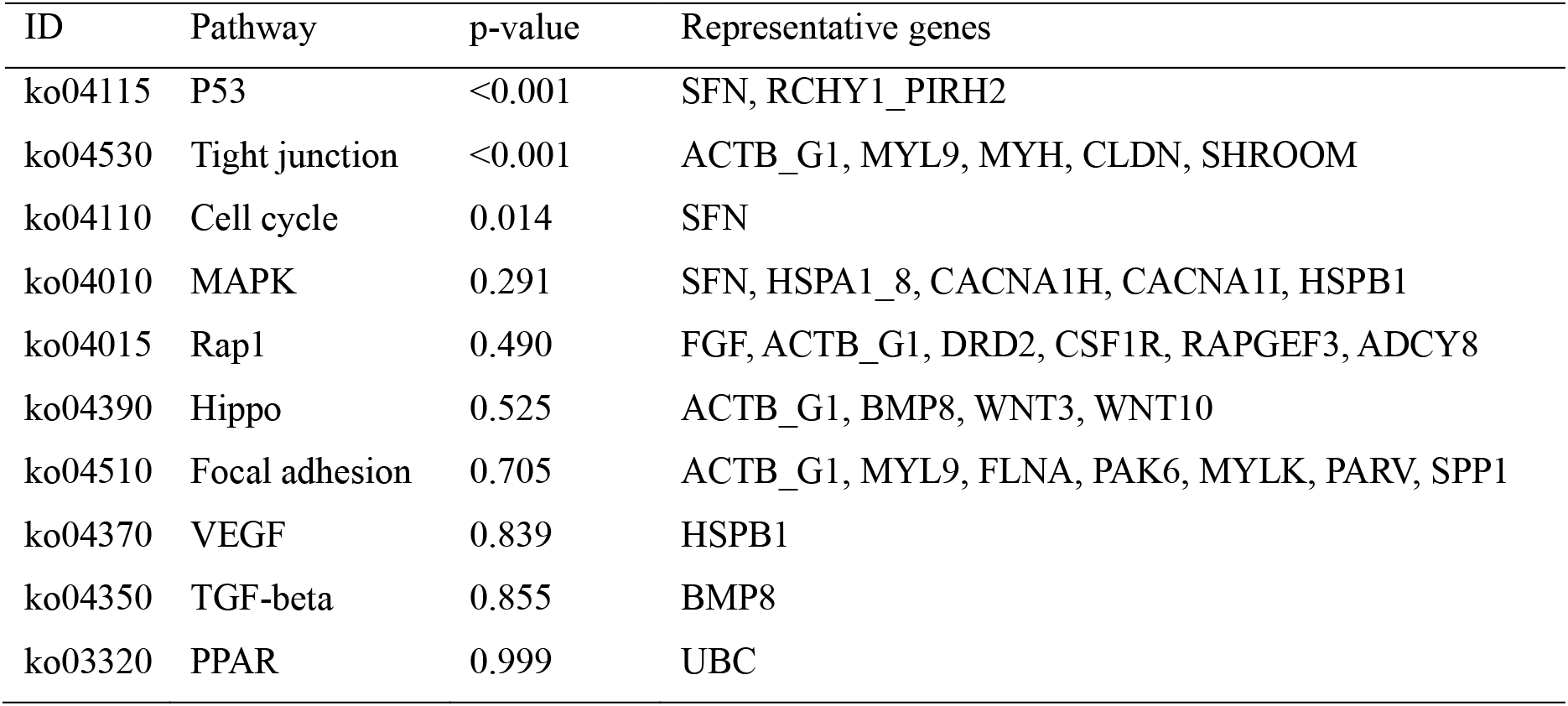
Summary of functional annotations of assembled unigenes with public protein databases.

In our study, 70,833 unigenes were assigned to 56 sub-categories of GO terms belonging to the following three main categories: biological process (BP), cellular component (CC), and molecular function (MF). These main categories included 26, 20, and 10 sub-term categories, respectively (Fig. 2). The most enriched GO terms were related to cellular and metabolic processes (BP), cell and organelle (CC), and binding and catalytic activity (MF). Importantly, we found some GO terms (level 4) in the BP category related to skin appendage development and differentiation, including (positive/negative) regulation of cell differentiation, regulation of cell proliferation, reproductive structure development, cell development, and ovarian follicle cell development.

**Figure 2.**
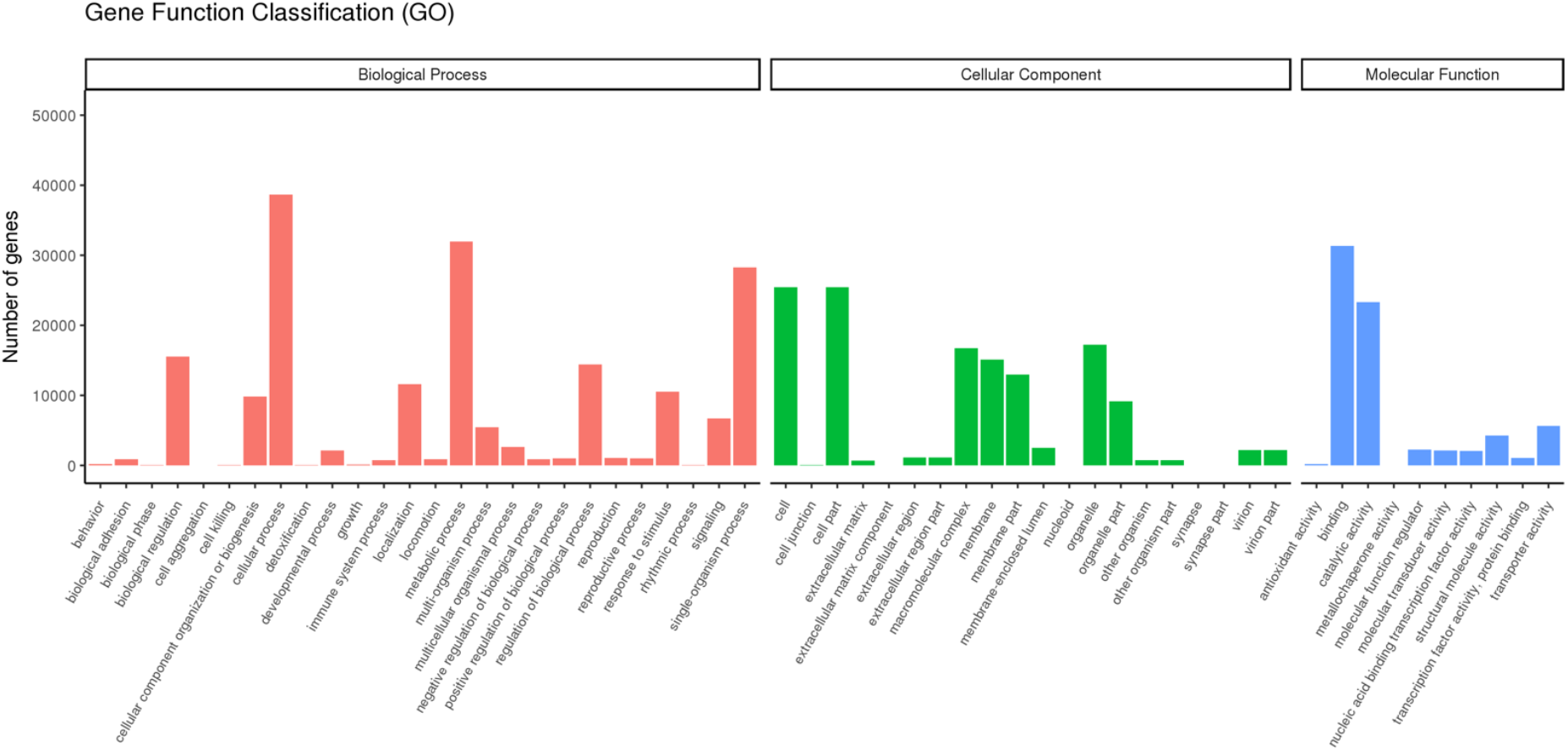
Gene Ontology classification of assembled unigenes in *Atelerix albiventris*.

Furthermore, 36,283 unigenes were mapped to 232 biological signaling pathways in KEGG database. Of which 15, 30, 22, 98, and 72 unique KEGG pathways represented in five KO hierarchies (Fig. 3). The most genes were annotated in translation pathways (5743 unigenes), next by signal transduction (5551 unigenes). In addition, there were some unigenes and pathways were related to skin appendage development and differentiation, including Rap1 (814 unigenes), Wnt (355 unigenes), Hippo (626 unigenes), TGF-b (206 unigenes), and Notch (171 unigenes) signaling pathways.

**Figure 3.**
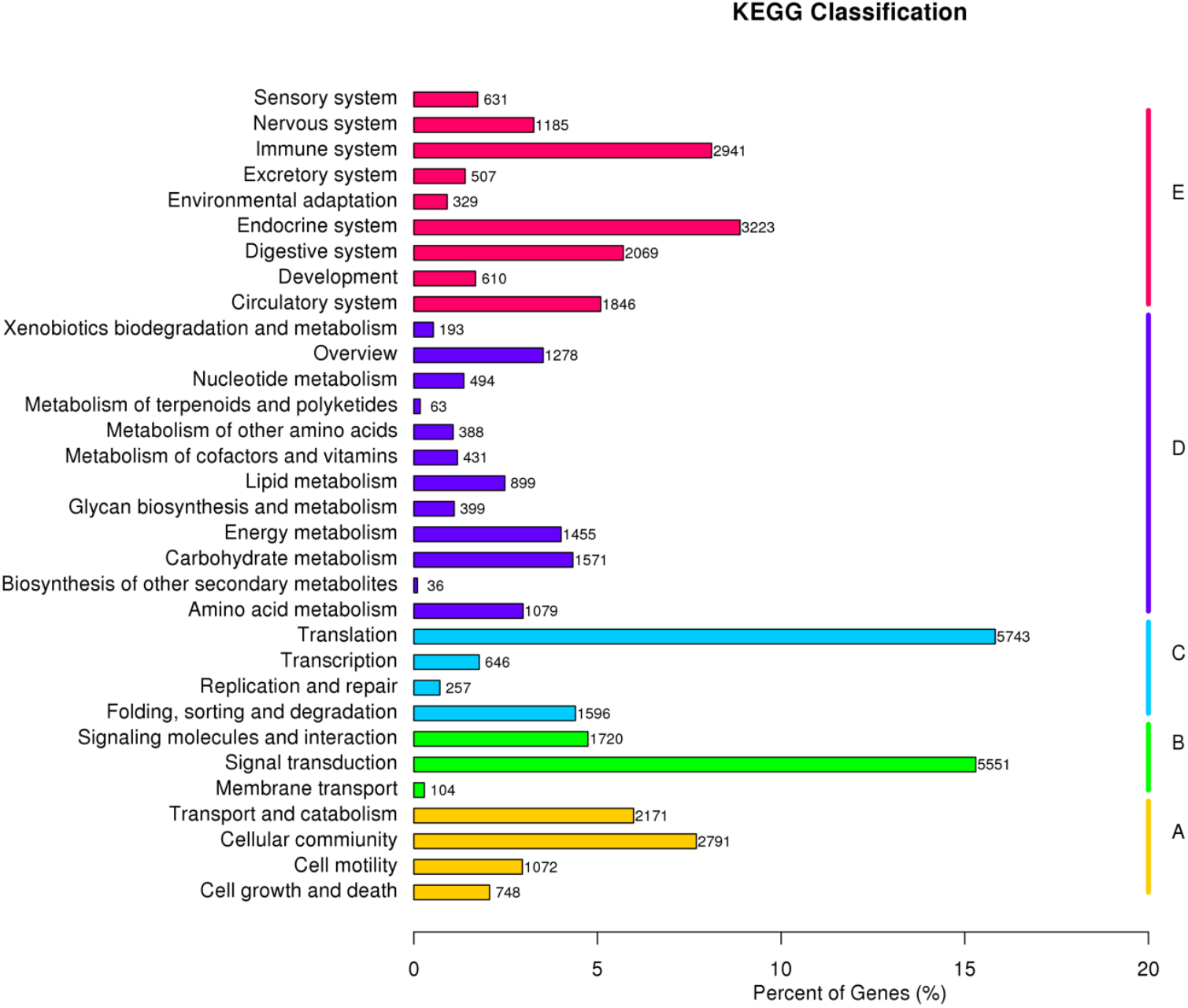
KEGG classification of assembled unigenes in *Atelerix albiventris*. (A) cellular processes; (B) environmental information processing; (C) genetic information processing; (D) metabolism, (E) organismal systems.

### Analysis of differentially expressed genes in skin appendage tissues

Analysis of DEGs in skin appendage tissues finally identified 3,598 DEGs in the three developmental stages, and 816 DEGs shared by the two types of tissues (Fig. 4a-c). As shown in the Venn diagram (Fig. 4d), more DEGs were found in tissues with spines (HH1Y and HS1Y) than in tissues without spines (HH1N and HS1N).

**Figure 4.**
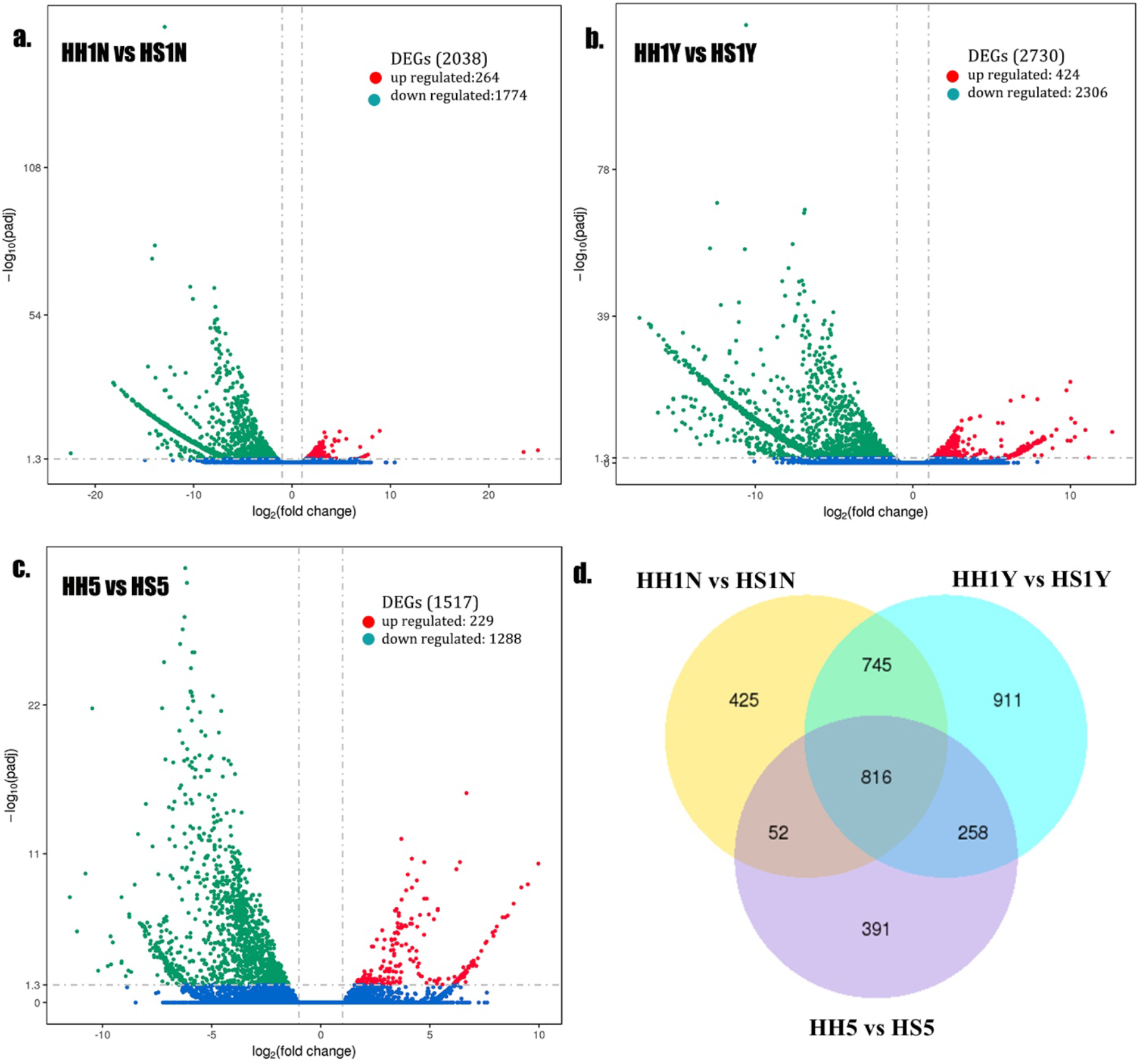
Differential expression analysis of hair- and spine-type tissues in *Atelerix albiventris*. Volcano plot of DEGs for (a) ‘HH1N vs HS1N’, (b) ‘HH1Y vs HS1Y’, (c) ‘HH5 vs HS5’. (d) Venn diagram showing co-DEGs among different tissues in *Atelerix albiventris.* DEGs, differentially expressed genes.

To validate expression patterns indicated by the transcriptome data, 11 differentiation-related DEGs were selected for RT-qPCR analysis including *KRT2, LEF1, RSPO2, ARRB, TGFB2, GRB2, SFN, TCF7, CTNNB1, HOXC13*, and *WIF1* (Supplementary Table 2). Expression trends determined by RT-qPCR significantly correlated with the RNA-Seq data (Fig. 5a). Expression of these 11 genes at the three developmental stages was also analyzed, revealing expression patterns similar to those determined by RNA-sequencing (Fig. 5b). Expression of *LEF1, TGFB2, SFN*, and *WIF1* at stages I–III increased significantly. However, expression of *KRT1* and *RSPO2* gradually decreased. Overall, both approaches confirmed the observed DEG trend patterns, indicating the accuracy of the transcriptome data and *de novo* RNA-Seq data.

**Figure 5.**
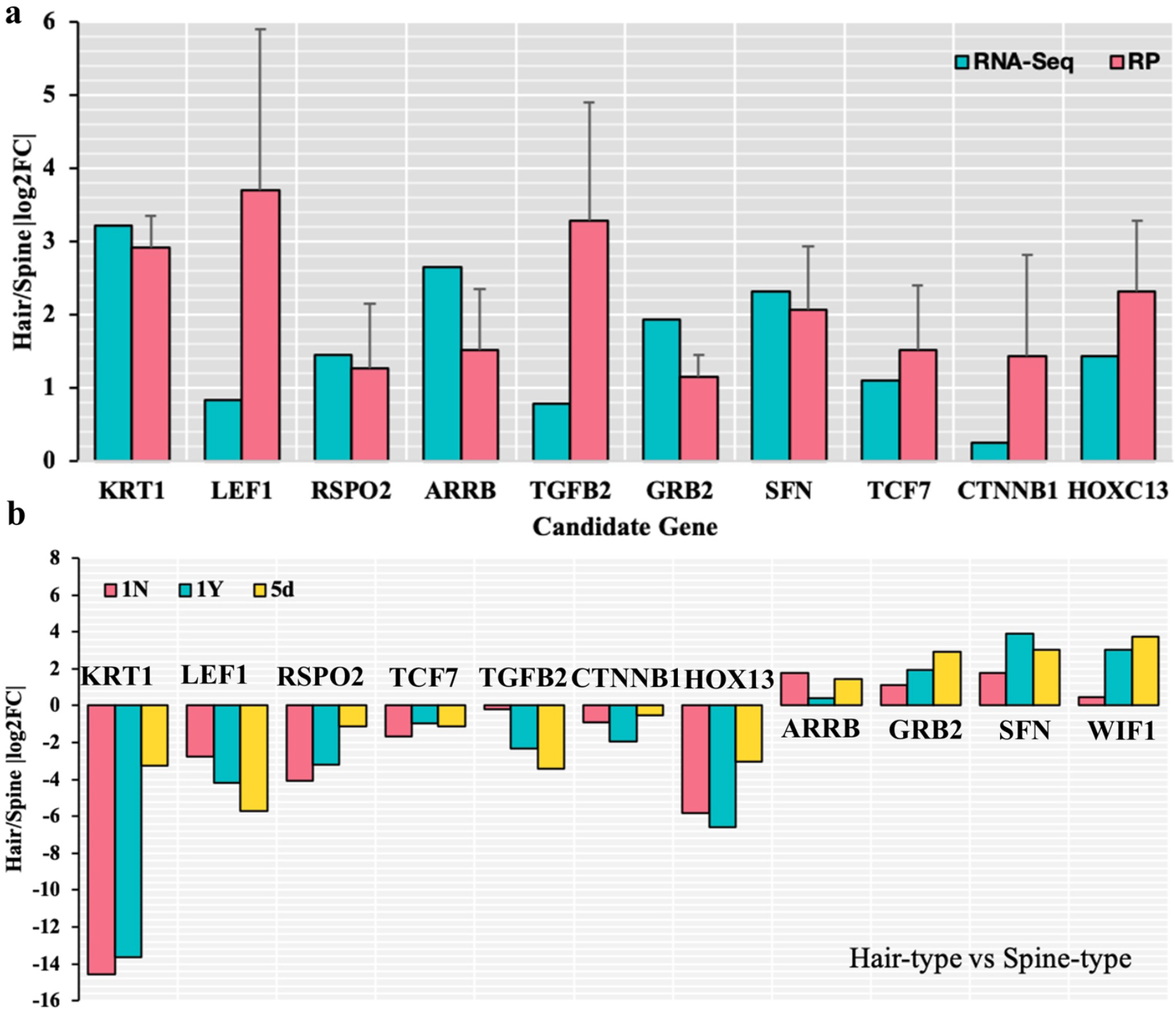
Quantitative real-time PCR analysis. (a) qPCR confirmation of 10 DEGs identified by RNA-seq in stage III tissues; (b) Relative expression levels of 10 DEGs in tissues at 3 developmental stages. Bars represent the relative expression levels of unigenes in stage III tissues normalized with respect to the internal control GAPDH. Error bars represent the standard error of three biological replicates. DEGs, differentially expressed genes.

### Differentially expressed genes related to skin appendage development

In order to further understand the spine development mechanism of the hedgehog, we conducted DEG analysis before and after spine development. In total, 28 DEGs were identified between HS1N and HS1Y, among them, 11 genes were downregulated, and the remaining 17 genes were upregulated (Supplementary Table 3). Based on GO and KEGG annotation, we found one downregulated gene of unknown open reading frame (LOC103122410) related to cell differentiation and multi-cellular organismal development in the BP category, and one downregulated keratin-associated protein like gene (LOC103118355) related to keratin filament.

In order to identify genes that activate the development of hedgehog abdominal hair, we analyzed DEGs before and after the occurrence of abdominal hair. A total 82 DEGs were identified between HH1N and HH1Y, of which 62 genes were downregulated, and the remaining 20 genes were upregulated (Supplementary Table 4). Notably, LOC103122410 was also present in these DEGs; in addition, keratin gene *KRT2* was highly expressed in HH1Y. We speculate that these genes play an important role in activating the development of hedgehog abdominal hair and spines.

### Differentially expressed genes related to skin appendage differentiation

We identified 1,517 DEGs by comparing the gene expression profiles of spine- (HS5) and hair-type (HH5) tissues in *A. albiventris* five days after birth. We screened 2 genes related to cell differentiation (*NOTCH* and keratin-associated protein (*KRTAP*) 9-2), 18 genes related to cell proliferation (*TGFB1, FGF, KRT2, TCHH, BMP3, BMP8, APOA1,CCDC85A, INHBB, MICAL2, VSIG8, FCN, GDF5, DMD, GDNF, FAT1, CSF*, and *GDF15*), and many keratin-related genes (*KRT1, KRT2, KRTAP*, etc). These genes might be indicative of hedgehog transcriptome involvement in spine and hair differentiation.

Based on annotation of the *A. albiventirs* transcripts with the GO database, the DEGs were significantly enriched in keratin-related terms. Keratin filament is an important component of skin appendages; the main DEGs involved in this process included *KRT1, KRT2, MYH, DES, BMP8, SHH*, and several *KRTAP* genes (Fig. 6a, b). In addition, *CSTA* was involved in the biological process of keratinocyte differentiation. In a directed acyclic graph diagram associated with this term, cell differentiation, epithelial cell differentiation, and epidermis development were enriched, and the main DEGs involved in this process included *KRT1, KRT2, MBP8, FGF, Wnt10*, and *MYH* (Fig. 6b). The only significantly enriched KEGG pathways among all DEGs comparing HH5 and HS5 tissues were tight junction and p53 signaling pathways (Fig. 6c; Table 2). In addition, eight pathways related to skin appendage development and differentiation were enriched, including cell cycle, MAPK, Rap1, Hippo, VEGF, TGF-beta, and PPAR signaling pathways. It is noteworthy that *SFN, BMP8, Wnt3, Wnt10, MYH*, and *SFN* were found in both annotation results, suggesting they are closely related to differentiation of keratinocytes and epidermal cells.

**Table 2.**
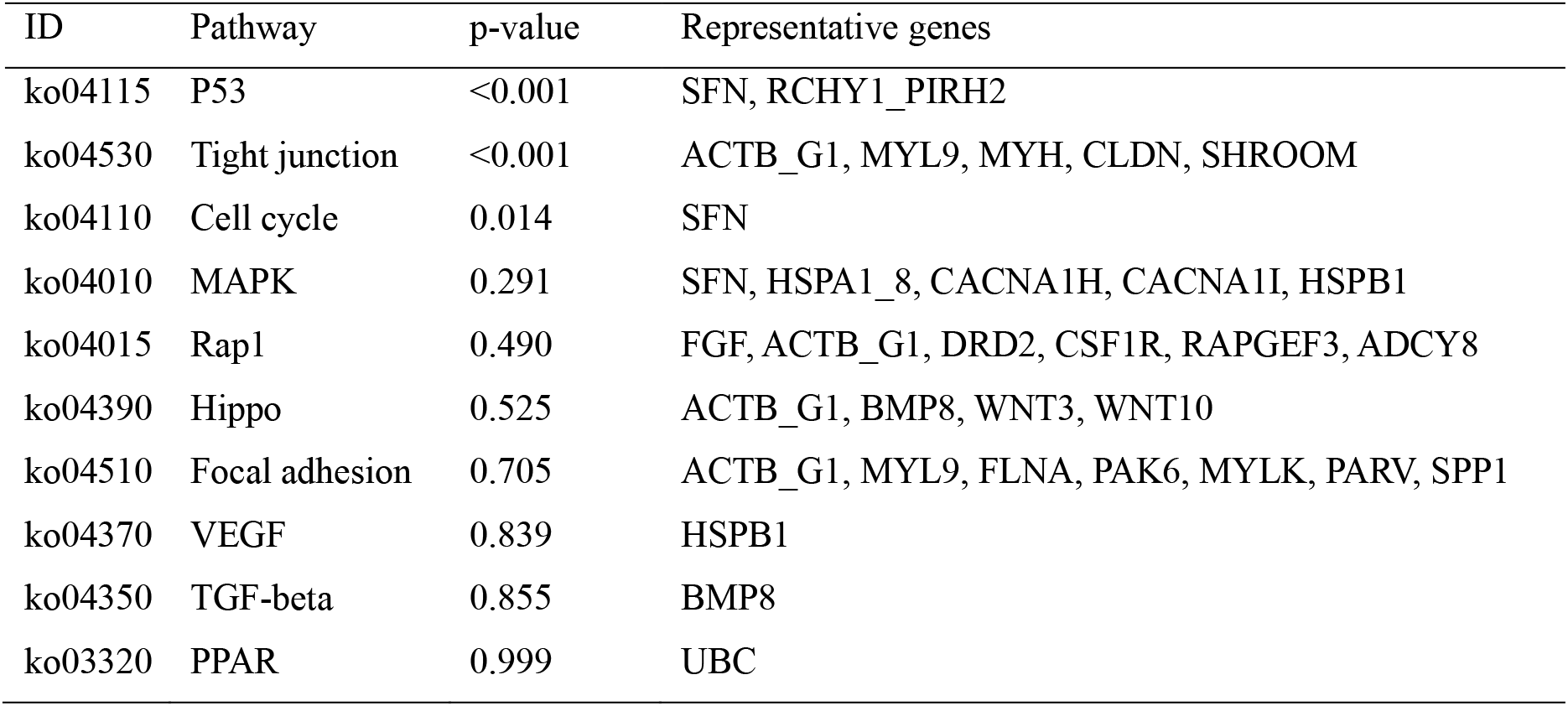
Candidate pathways and genes related to development and differentiation of spine and hair.

**Figure 6.**
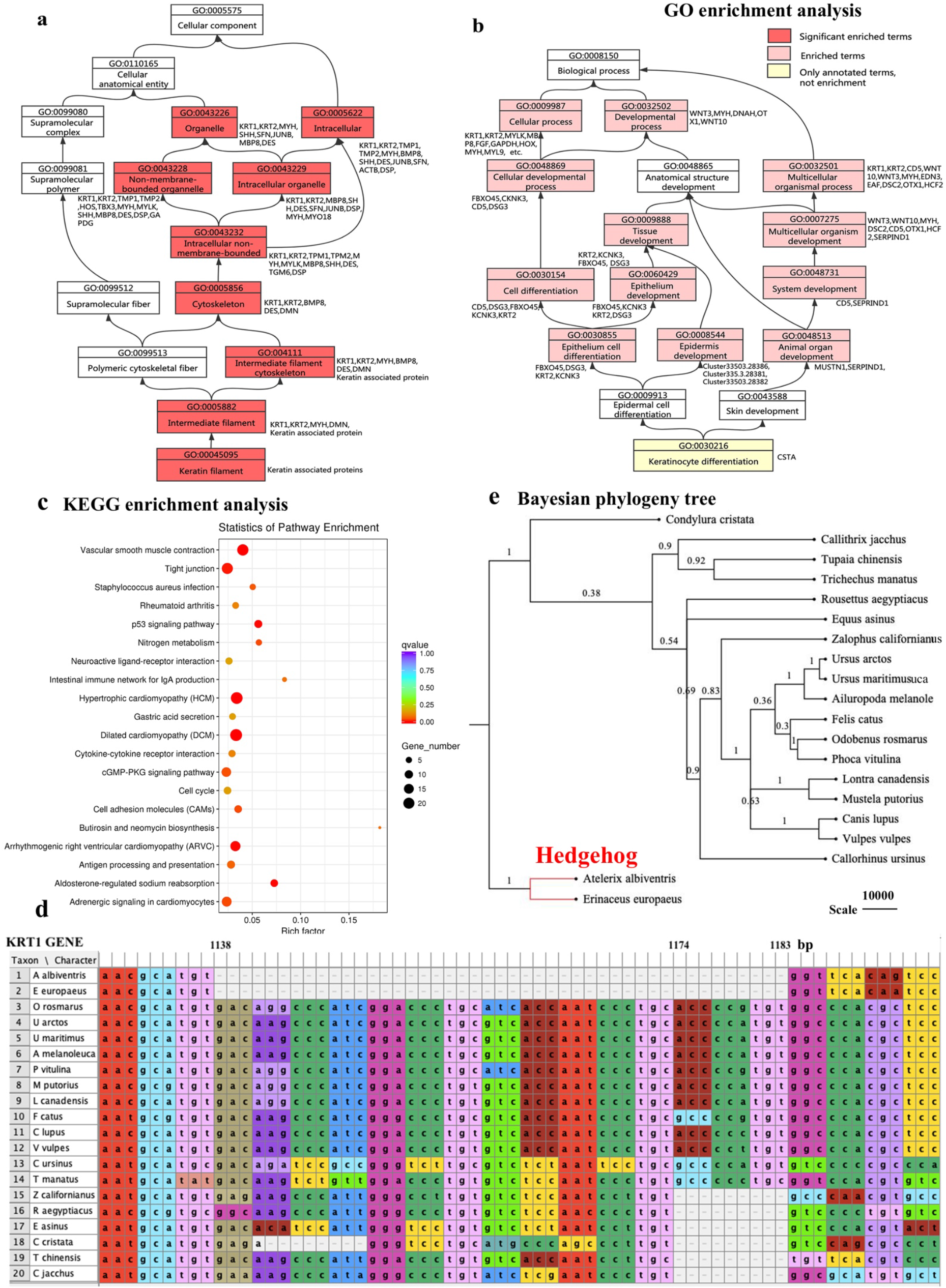
KEGG and GO enrichment analysis of DEGs related to differentiation of KRT1 gene. (a,b) Directed acyclic graph related to skin appendage development and differentiation in GO enrichment analysis. (c) Dot plots of enriched KEGG pathways. (d) Multiple sequence alignment of *KRT1* gene from 20 species. (e) Bayesian phylogenetic tree with *KRT1* sequences from 20 species. DEGs, differentially expressed genes.

In our data, keratin-related genes were highly abundant and also significantly overexpressed in the dorsal spine-type tissues. In the hedgehog transcriptome, more than 3,000 transcripts were annotated to 6 keratin genes (*KRT1, KRT2, KRT5, KRT10, TCHP*, and *CHRNA1*) and approximately 20 keratin-associated proteins. Through alignment of *KRT1* gene sequences from 20 species, we found that the hedgehog *KRT1* gene sequence has a deletion of 45 bp compared with other species (Fig. 6d). The two hedgehog species were isolated into one clade in the bayesian phylogenetic tree (Fig. 6e), suggesting that *KRT1* may be one of the important regulatory genes for the development and differentiation of hedgehog skin appendages.

### Immune-related genes in epidermis of hedgehog

In this study, we also found many immune-related genes comparing HH5 and HS5. After KEGG enrichment analysis, we found no significant enrichment of immune-related pathways in spine-type tissues, and fewer immune-related pathways and genes than in hair-type tissues. Hair-type tissues exhibited 19 enriched immune-related pathways, three of which were significantly enriched, and 47 transcripts annotated with 32 immune-related genes (Fig. 7a,b). In addition, we observed similar GO enrichment analysis results. The number of terms and genes enriched in hair-type tissues was significantly higher than in spine-type tissues, of which two of the seven terms were significantly enriched, and 25 transcripts were annotated with 11 immune-related genes (Fig. 7c,d).

**Figure 7.**
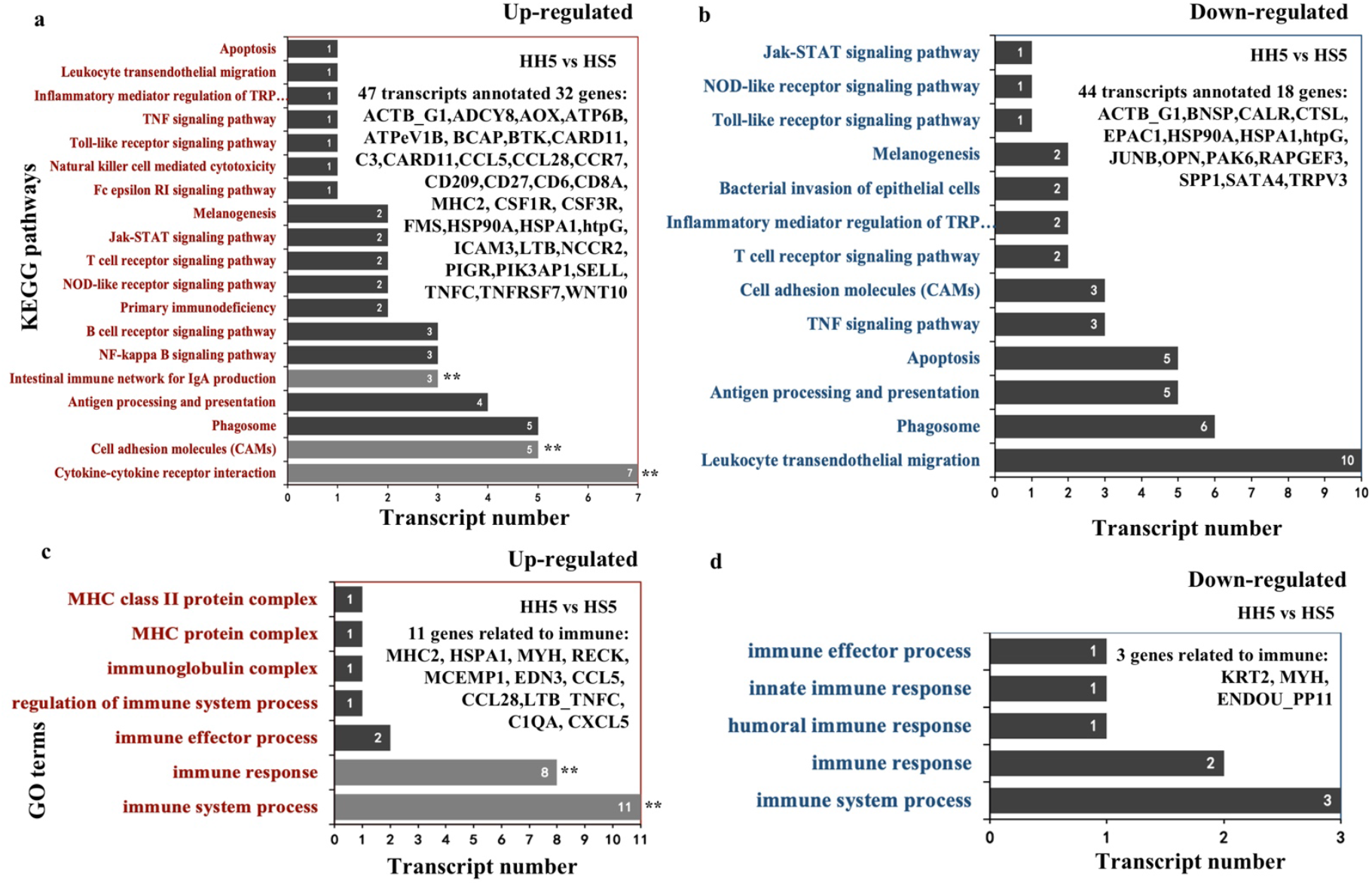
KEGG and GO enrichment analysis of DEGs related to immunity. (a,b) up- and downregulated KEGG signaling pathways and key genes related to immunity. (c,d) up- and downregulated GO terms and key genes related to immunity. DEGs, differentially expressed genes.

### Discussion

The genetic basis of morphological variation, both within and between species, provides a major topic in evolutionary biology. In mammals, the development of skin appendages such as hair, tooth, and scale involves complex interactions between the epidermis and the underlying mesenchyme as part of an established hierarchical morphogenetic process^21^. Specifically, mammals develop a coat containing many distinct types of hair. Such diversity is associated with molecular and signaling pathways that drive formation and induction in a specific spatial and temporal manner^22^. Reciprocal interactions between epithelial and mesenchymal tissues constitute a central mechanism that determines the location, size, and shape of organs^23^.

In the current study, we aimed to explore the genetic basis for hedgehog skin appendage differentiation and development and the resulting expression of the spine trait. We identified 328,576 unigenes in our transcriptome, all of which were annotated in 7 databases (Table 1). Taken together, our *de novo* assemblies revealed higher quality compared with previous studies^24,25^ that formed the foundation of all our subsequent analyses. According to GO and KEGG annotation analysis, a total of 70,833 and 36,238 unigenes were mapped to 56 sub-categories and 232 biological pathways, respectively (Fig. 2,3). We identified some of the key pathways involved in skin development; these annotations may provide a valuable resource for further understanding the specific functions and pathways in *A. albiventris*.

Newborn hedgehogs begin to develop hair/spines approximately two hours after birth. We found 6 shared genes (*APOE, COX2, COX3, FCN, RP-L11e, SH3GL*) through analysis of DEGs before and after hair/spine development, indicating that these genes are essential regulatory genes in the development of skin appendages, whether hair or spine. Further, we compared the gene expression profiles of spine- and hair-type tissues to systematically assess key regulatory genes for spine and hair differentiation in *A. albiventris*. *SFN*, upregulated in spine-type tissues, is a regulator of mitotic translation that interacts with a variety of translation and initiation factors^26^, is enriched in the p53 signaling pathway, and has a role in keratinocyte differentiation and skin barrier establishment in the BP category (GO). *SFN* plays an important role in maintaining hair follicle development, especially affecting the formation of hair shaft structure^27^. Bu *et al*. (2014) determined that SFN gene and protein were significantly highly expressed in the thicker, longer, and harder skin of wool, indicating that *SFN* was involved in the regulation of wool character development^28^. In addition, we identified *FGF* as the upregulated DEG enriched in the MAPK and Rap1 signaling pathways. These pathways often act together by forming signaling loops during organogenesis^29^ and induce the most fundamental biological processes, such as the formation of periodic patterns. Hebert *et al.* (1994) demonstrated that *FGF* plays an important role in regulation of the hair cycle growth, and functions as an inhibitor of hair elongation by promoting progression from anagen stage, the growth phase of the hair follicle, to catagen stage, the apoptosis-induced regression phase^16^. Hence, we believe that *SFN* and *FGF* may be important regulators that affect spine development and differentiation in *A. albiventris*.

We also compared *KRT1* gene sequences of 20 mammals with that of the hedgehog, in which we identified a 45-bp deletion in the latter. Keratins, the major structural proteins of epithelia, are a diverse group of cytoskeletal scaffolding proteins that form intermediate filament networks and provide structural support to keratinocytes that maintain skin integrity^30^. In general, *KRT1* is highly conserved in mammals, however, molecular defects in keratin intermediate filament-related genes can cause keratinocyte and tissue-specific fragility, accounting for a large number of genetic disorders in skin and its appendages^31,32^. Therefore, whether the deletion phenomenon in hedgehog *KRT1* has special significance for spine development and differentiation in the hedgehog warrants further research and verification.

Lastly, we analyzed immune-related genes in the skin transcriptome of *A. albiventris*. We found that hair-type skin had more immune-related genes than spine-type skin. Choo *et al.* (2016) found that interferon epsilon (*IFNE*), which was exclusively expressed in epithelial cells and was important to mucosal immunity, was pseudogenized in pangolins. They proposed that scale development provided protection against injuries or stress and reduced pangolin vulnerability to infection, thus protection by scales on the pangolin body compensated for the low immunity of this species to a certain extent^33^. From our current data, we speculate that there may be significant differences in the immune function of skin with different appendage types, and that the occurrence of spines may be an innovative physical protection for dorsal skin with relatively low immunity.

### Conclusions

In the current study, we conducted a comprehensive transcriptome analysis of *A. albiventris* to explore the genetic basis for the growth of skin appendages. Transcriptome analysis provided a rich list of unigenes expressed in hair- and spine-type tissues at three different developmental stages, of which 328,576 unigenes and 3,598 DEGs were identified. Candidate genes were identified that are likely involved in the regulation of hair and spine growth and differentiation. The knowledge acquired in this study of the molecular and signaling pathways related to hair and spine expression greatly contributes to the current genetic resources for the hedgehog and mammalian species that harbor shaggy appendages, as well as traits that could be potentially exploited for curing skin diseases of other animals, even humans.

## Methods

### Ethics statement

All animal procedures in the study were approved by the ethics committee for animal experiments at the Guangdong Institute of Applied Biological Resources (reference number G2ABR20170523) and followed basic principles. We confirm that all methods were performed in accordance with relevant guidelines and regulations.

### Biological samples

According to observations of the growth characteristics of hedgehogs, hairs and spines on the back and abdomen begin to appear approximately two hours after birth; therefore, we obtained 8 hedgehogs at 3 different stages of appendage development from a commercial animal farm (Dongguan City, China). Hedgehog hair (HH) and spine (HS) tissues were collected from 2 specimens within 2 h of birth (Stage I, HH1N/HS1N), 3 specimens after 2 h but within the first birth day (Stage II, HH1Y/HS1Y), and 3 specimens 5 days after birth (Stage III, HH5/HS5). Prior to skin sampling, follow the standard operating procedures for euthanasia, conduct euthanasia for experimental animals, minimize the pain of small animals, do not affect the results of animal experiments, and shorten the time of death as far as possible. Sixteen skin tissue samples representing the two types of appendages (abdomen hair-type and dorsal spine-type) were rapidly excised, immediately snap-frozen on dry ice, and stored at –80 °C until RNA extraction.

### RNA extraction and RNA-seq

Total RNA from each tissue sample was extracted using the RNeasy Kit (Qiagen, Hilden, Germany). RNA purity was determined using a NanoPhotometer^®^ spectrophotometer (Implen, Inc., Westlake Village, CA, USA). All unigenes were annotated using the basic local alignment search tool (BLASTX), considering hits with e-values of 1E-5 against seven databases: NCBI non-redundant protein sequences (Nr); NCBI non-redundant nucleotide sequences (Nt); Protein family (Pfam) database; Clusters of Orthologous Groups of Proteins (KOG/COG) database; Swiss-Prot (a manually annotated and reviewed protein sequence database); KEGG Ortholog (KO) database; Gene Ontology (GO) database. Differentially expressed genes (DEGs) were identified by comparing gene expression levels between samples (or sample groups). Bowtie2^34^ was used to align clean reads with all unigenes and RSEM^35^ was used to calculated gene expression levels for each sample. Differential gene expression was analyzed using DESeq270 V1.16.1 in RStudio71 V1.0.143 running R72 V3.4.1. All sequencing was conducted by Novogene (Beijing, China).

### Quantitative real-time reverse-transcription PCR (RT-qPCR)

RNA was extracted from skin appendage tissue samples using TRIzol reagent (Invitrogen, Carslbad, CA, USA). Then, cDNA was synthesized using the Toyobo reverse-transcription kit (Toyobo, Osaka, Japan). Eleven genes related to hair and/or spine development were selected for analysis based on functional annotation data, and fluorescent qPCR primers were designed accordingly (Supplementary Table 2). RT-qPCR was performed in a 20-μL reaction volume, with four technical replicates for each sample, using the TransStart^®^ Top Green qPCR SuperMix kit (TransGen Biotech, Beijing, China). Relative gene expression levels were analyzed using the 2^(-DDCT) method (Bustin *et al*. 2010). PCR conditions were as follows: pre-denaturation at 95 °C for 10 min; 40 cycles of 15 s at 95 °C (denaturation), 30 s at 58 °C (annealing), and 20 s at 72 °C (extension); and a final melting curve stage from 60-95 °C to verify the specificity of the amplicons.

### Phylogenetic analysis

*KRT1* sequence authenticity was verified by BLAST search in GenBank (Supplementary Table 5). Sequences were edited using MAFFT v 6.81b^36^ and Mesquite^37^. MRMODELTEST v.2.3^38^ was used to select the best-fit model of nucleotide substitution under the Akaike information criterion (AIC)^39^. Bayesian inference of phylogeny was performed using BEAST v 1.6.1^40^ with default settings except for GRT+I+G model. An uncorrelated relaxed clock fixed to lognormal distribution was employed as the site model using a Yule speciation tree prior sampled every 10,000th generation for 100 million generations. Effective sample seizes (ESS) were verified using Tracer v1.5 and a consensus tree was constructed in TreeAnnotator v1.6.1 with 20% burn-in. For the concatenated dataset, all parameters were estimated independently for each partition and displayed using FigTree v1.4.2. (http://www.geospiza.com/finchtv).

## Abbreviations

*A. albiventris*: *Atelerix albiventris*
Hiseq: high-throughput sequencing
PCA: principal component analysis
KRT: keratin
FGF: fibroblast growth factor
SHH: sonic hedgehog
BMP: bone morphogenetic protein
DEGs: differentially expressed genes
RT-qPCR: quantitative real time polymerase chain reaction
SOM: self-organizing feature map

## Declarations

### Ethics approval and consent to participate

All animal procedures were approved by the ethics committee for animal experiments at the Guangdong Institute of Applied Biological Resources (reference number G2ABR20170523) and followed basic principles.

### Consent for publication

Not applicable

### Availability of data and materials

Data analyzed in the current study are included within the article and its supplementary material. All unigene sequences from *A. albiventris* have been deposited in the GenBank Sequence Read Archive (SRA) under accession number PRJNA561241 for SUB6195278. We have uploaded supplemental material to figshare via the GSA Portal. Supplementary Table 1 contains the summary of sequencing statistics for the transcriptomes; Supplementary Table 2 contains the summary of primer information used in real-time PCR analysis; Supplementary Table 3 contains the differentially expressed genes between HS1N and HS1Y; Supplementary Table 4 contains the differentially expressed genes between HH1N and HH1Y; Supplementary Table 5 contains the species name and NCBI serial numbers of 19 additional species for phylogenetic analysis.

### Authors’ contributions

H. Li and J. Chen designed research; H. Li analyzed data and wrote the report; B. Yang developed software necessary to perform and record experiments; L. Li, X. Zhang, H. Ahmad, and H. Jiang provided expertise and advice on computational analysis; all authors edited the report.

## Acknowledgments

We gratefully acknowledge the assistance of Professor Gang Li in the design of the experiments. We thank Wiley publishing (http://wileyeditingservices.com) for editing the draft of this manuscript.

## Supplementary tables

**Supplementary Table 1 Sample IDs for transcriptome sequencing**

**Supplementary Table 2 Summary of primers for real-time PCR analysis**

**Supplementary Table 3 Differentially expressed genes between HS1N and HS1Y**

**Supplementary Table 4 Differentially expressed genes between HH1N and HH1Y**

**Supplementary Table 5 NCBI accession numbers of *KRT1* sequences from 19 species**

