## Supplementary table 1-5 for "Transcriptome analysis reveals the genetic basis underlying the development of skin appendages and immunity in hedgehog (*Atelerix albiventris*)"

**A****bstract**

The expression of hair features is an evolutionary adaptation resulting from interactions between many organisms and their environment. Elucidation of the mechanisms that underlie the expression of such traits is a topic in evolutionary biology research. Therefore, we assessed the *de novo* transcriptome of *Atelerix albiventris* at three developmental stages and compared gene expression profiles between abdomen hair and dorsal spine tissues. We identified 328,576 unigenes in our transcriptome, among which 3,598 were differentially expressed between hair- and spine-type tissues. Dorsal and abdomen skin tissues 5 days after birth were compared and the resulting differentially expressed genes were mainly enriched in keratin filament, epithelium cell differentiation, and epidermis development based on GO enrichment analysis, and tight junction, p53, and cell cycle signaling pathways based on KEGG enrichment analysis. Expression variations of *MBP8, SFN, Wnt10, KRT1*, and *KRT2* may be the main factors regulating hair and spine differentiation for the hedgehog. Strikingly, DEGs in hair-type tissues were also significantly enriched in immune-related terms and pathways with hair-type tissues exhibiting more upregulated immune genes than spine-type tissues. Thus, we propose that spine development was an adaptation that provided protection against injuries or stress and reduced hedgehog vulnerability to infection.

**Keywords:** hedgehog, skin appendage, adaptive evolution, RNA-Seq, molecular biology

Supplementary table 1 Sample IDs for transcriptome sequencing

| **Sample ID** | **RNA integrity number （RIN）** | **Concen.**  **(ng/ul)** | **Raw Reads** | **Clean reads** | **Clean bases**  **(G)** | **Error**  **(%)** | **Q20**  **(%)** | **Q30**  **(%)** | **GC**  **(%)** |
| --- | --- | --- | --- | --- | --- | --- | --- | --- | --- |
| HH1_N1 | 9.3 | 414 | 84746676 | 82076208 | 12.3 | 0.03 | 96.85 | 92.17 | 58.14 |
| HH1_N2 | 7.6 | 252 | 91823824 | 89358204 | 13.4 | 0.03 | 97.2 | 93.01 | 58.02 |
| HS1_N1 | 8.4 | 1455 | 87533090 | 84776044 | 12.72 | 0.03 | 96.44 | 91.23 | 58.13 |
| HS1_N2 | 7.8 | 1986 | 80827066 | 78473126 | 11.77 | 0.03 | 95.92 | 90.09 | 57 |
| HH1_Y1 | 8.6 | 324 | 78351662 | 76208670 | 11.43 | 0.03 | 97.01 | 92.49 | 57.67 |
| HH1_Y2 | 8.4 | 416 | 83163428 | 80515340 | 12.08 | 0.03 | 97 | 92.5 | 58.74 |
| HH1_Y3 | 8.9 | 674 | 91357512 | 88846572 | 13.33 | 0.03 | 97.3 | 93.23 | 58.61 |
| HS1_Y1 | 8.9 | 1290 | 97436460 | 94259192 | 14.14 | 0.03 | 96.46 | 91.35 | 58.29 |
| HS1_Y2 | 8.6 | 1350 | 92684804 | 89648842 | 13.45 | 0.03 | 96.4 | 91.22 | 58.46 |
| HS1_Y3 | 6.6 | 90 | 86012194 | 84258058 | 12.64 | 0.03 | 97.05 | 92.27 | 49.4 |
| HH5_1 | 8.3 | 134 | 81247180 | 78253286 | 11.74 | 0.03 | 97.3 | 93.26 | 59.02 |
| HH5_2 | 9.5 | 42 | 92765874 | 90002040 | 13.5 | 0.03 | 96.84 | 92.14 | 54.7 |
| HH5_3 | 7.8 | 338 | 87389764 | 84746764 | 12.71 | 0.03 | 97.1 | 92.81 | 58.38 |
| HS5_1 | 8.8 | 204 | 100102332 | 97325590 | 14.6 | 0.03 | 96.42 | 91.29 | 58.04 |
| HS5_2 | 7.1 | 204 | 98119576 | 95197288 | 14.28 | 0.03 | 96.36 | 91.15 | 56.88 |
| HS5_3 | 6.4 | 78 | 87681000 | 84955022 | 12.74 | 0.03 | 96.35 | 91.06 | 55.88 |

Supplementary table 2 Summary of primers of real-time PCR assay

| Gene ID | Symbol | Primer-F (5'-3') | Primer-R (3'-5') |
| --- | --- | --- | --- |
| Cluster-33503.52810 | ARRB | AAAACAGAAAGGGAAACG | AGGAGGTGGGCGAAGA |
| XM_007527837.2 | GAPDH | ACTCCACTCACGGCAAAT | GTACTCGGCACCAGCATC |
| Cluster-33503.134689 | GRB2 | GGGGTTTGATGCGGATAT | GCCACCGAAAGTGACGAG |
| Cluster-33503.28078 | HOXC13 | GTGACCCTGGAGCAAA | CTATCTATTAGTGGGACTTGG |
| Cluster-33503.122683 | KRT1 | AGCCGCAGTTGCCCACA | CCAGCACGATGCCTTACA |
| Cluster-33503.128985 | LEF1 | TTTCCGTAGTTGTCCCG | TGAATGCGTTCATGCTGT |
| Cluster-33503.47741 | RSPO2 | CCGGAGACCCTGGAGTTGT | GGACCGCCAGAGGCAATT |
| Cluster-33503.95190 | SFN | CTCACTTCACAGAGCCTTTC | GCTGCTGCGAGACAACCT |
| Cluster-33503.30444 | TCF7 | CAAAGTGATTGCCGAGTG | TTGTCCCGTGCTGACC |
| Cluster-33503.107807 | TCNNB1 | CTTCTGGGCTACGATGAC | CAACTCTGCTTCCTGGTG |
| Cluster-33503.38152 | TGFB2 | GAACCCGACTGTGCTGA | TGCCTCCGTCCTCTTTA |
| Cluster-33503.47776 | WIF1 | AAGTTCGTCTGTAGCGTGAT | CCTTCTCCAATGTTCCCT |

Supplementary table 3 Differentially expressed genes between HS1N and HS1Y

| Gene ID | HS1N_readcount | HS1Y_readcount | log2FoldChange | p-value | padj |
| --- | --- | --- | --- | --- | --- |
| Cluster-33503.128444 | 0 | 86.08214993 | -21.625 | 9.31E-07 | 0.010868 |
| Cluster-33503.125788 | 0 | 63.24402852 | -21.197 | 1.52E-06 | 0.016849 |
| Cluster-33503.71896 | 0 | 30.45082855 | -20.179 | 4.74E-06 | 0.036887 |
| Cluster-33503.160189 | 0 | 43.12854788 | -7.8749 | 2.08E-06 | 0.020771 |
| Cluster-33503.115414 | 1.453397872 | 57.98866115 | -5.2885 | 2.29E-06 | 0.021883 |
| Cluster-33503.50680 | 21.81424054 | 336.6388061 | -3.9508 | 5.45E-10 | 1.64E-05 |
| Cluster-33503.127585 | 42.52858859 | 558.0876554 | -3.7159 | 2.27E-11 | 7.94E-07 |
| Cluster-33503.89984 | 96.11450522 | 896.5460164 | -3.2208 | 4.10E-06 | 0.034448 |
| Cluster-33503.121879 | 25.91251299 | 236.8442846 | -3.1921 | 7.28E-07 | 0.009559 |
| Cluster-82032.0 | 18.83532828 | 172.1416422 | -3.1884 | 9.18E-07 | 0.010868 |
| Cluster-33503.129745 | 160.6011264 | 1414.024237 | -3.1376 | 7.34E-09 | 0.000141 |
| Cluster-33503.40595 | 11872.35656 | 2226.207379 | 2.4149 | 1.77E-06 | 0.018626 |
| Cluster-33503.107453 | 4977.639861 | 896.6663599 | 2.4726 | 3.93E-06 | 0.034428 |
| Cluster-33503.30824 | 514.2493259 | 71.4815556 | 2.8474 | 5.04E-06 | 0.03779 |
| Cluster-33503.86800 | 9843.401602 | 1356.213765 | 2.8594 | 4.34E-06 | 0.035042 |
| Cluster-33503.64008 | 192.0790232 | 19.91437288 | 3.2642 | 2.88E-07 | 0.004321 |
| Cluster-33503.107454 | 7002.950394 | 673.0459194 | 3.3788 | 2.45E-09 | 5.72E-05 |
| Cluster-33503.84059 | 409.0755235 | 25.82116204 | 3.9892 | 3.70E-12 | 1.56E-07 |
| Cluster-33503.40525 | 90.36019879 | 4.296914424 | 4.4067 | 4.09E-07 | 0.005723 |
| Cluster-33503.69414 | 1973.58419 | 87.84111687 | 4.4903 | 7.56E-20 | 5.29E-15 |
| Cluster-33503.93070 | 30.37756328 | 0 | 7.4075 | 2.73E-06 | 0.024973 |
| Cluster-33503.71791 | 1155.372535 | 5.005001278 | 7.8257 | 2.25E-32 | 2.36E-27 |
| Cluster-33503.74871 | 2263.274482 | 6.711504128 | 8.3611 | 3.22E-39 | 6.77E-34 |
| Cluster-33503.93079 | 135.0359235 | 0 | 9.5605 | 1.45E-12 | 7.61E-08 |
| Cluster-33503.43551 | 188.7802277 | 0 | 24.007 | 2.92E-08 | 0.000472 |
| Cluster-33503.96104 | 289.6354178 | 0 | 24.591 | 1.34E-08 | 0.000234 |
| Cluster-33503.78173 | 398.7659056 | 0 | 25.028 | 7.37E-09 | 0.000141 |
| Cluster-33503.83755 | 1125.957687 | 0 | 26.444 | 9.97E-10 | 2.62E-05 |

Supplementary table 4 Differentially expressed genes between HH1N and HH1Y

| Gene ID | HH1N_readcount | HH1Y_readcount | log2FoldChange | pval | padj |
| --- | --- | --- | --- | --- | --- |
| Cluster-33503.110885 | 0 | 66.63694 | -21.368 | 1.25E-06 | 0.004152 |
| Cluster-24458.2 | 0 | 32.18077 | -20.354 | 3.92E-06 | 0.010497 |
| Cluster-33503.125440 | 0 | 55.892 | -20.339 | 3.96E-06 | 0.010497 |
| Cluster-33503.27650 | 0 | 31.01105 | -20.297 | 4.17E-06 | 0.010871 |
| Cluster-33503.71896 | 0 | 30.28986 | -20.271 | 4.29E-06 | 0.011009 |
| Cluster-23971.0 | 0 | 35.42226 | -7.6936 | 1.78E-06 | 0.005659 |
| Cluster-50926.0 | 0.901617 | 96.81909 | -6.71 | 1.04E-08 | 6.86E-05 |
| Cluster-33503.28873 | 1.381753 | 62.27404 | -5.4875 | 3.64E-06 | 0.009968 |
| Cluster-33503.115414 | 6.575269 | 120.4126 | -4.202 | 5.87E-10 | 5.19E-06 |
| Cluster-33503.52438 | 22.75327 | 396.6611 | -4.1295 | 1.07E-12 | 2.43E-08 |
| Cluster-33503.45503 | 6.967422 | 111.498 | -4.001 | 3.33E-09 | 2.52E-05 |
| Cluster-33503.50611 | 18.53091 | 272.9855 | -3.88 | 2.69E-13 | 7.12E-09 |
| Cluster-27220.0 | 3.606467 | 53.88451 | -3.878 | 2.76E-06 | 0.007966 |
| Cluster-33503.93776 | 12.19027 | 163.9718 | -3.7533 | 8.46E-12 | 1.22E-07 |
| Cluster-33503.50604 | 8.349175 | 99.9293 | -3.5813 | 6.44E-08 | 0.000301 |
| Cluster-33503.50610 | 18.13876 | 206.5931 | -3.5106 | 5.67E-10 | 5.19E-06 |
| Cluster-33503.48696 | 6.428629 | 71.85253 | -3.4766 | 1.35E-06 | 0.004382 |
| Cluster-33503.50609 | 238.1785 | 2496.305 | -3.3895 | 7.59E-14 | 2.41E-09 |
| Cluster-33503.63866 | 11.34731 | 114.1013 | -3.3365 | 1.05E-07 | 0.000453 |
| Cluster-33503.50605 | 102.2464 | 946.8086 | -3.2117 | 2.29E-11 | 2.80E-07 |
| Cluster-33503.50612 | 53.87748 | 491.173 | -3.1887 | 2.24E-10 | 2.54E-06 |
| Cluster-33503.50603 | 127.7263 | 1118.064 | -3.13 | 2.65E-12 | 4.22E-08 |
| Cluster-33503.142390 | 14.18035 | 109.9152 | -2.9497 | 1.15E-05 | 0.025082 |
| Cluster-33503.46837 | 15.73808 | 116.9022 | -2.8927 | 1.83E-06 | 0.005715 |
| Cluster-33503.38566 | 49.4976 | 350.2752 | -2.8215 | 2.98E-10 | 3.16E-06 |
| Cluster-33503.99443 | 30.35835 | 203.1055 | -2.7441 | 1.26E-07 | 0.000526 |
| Cluster-33503.121879 | 55.20058 | 369.5113 | -2.7422 | 1.25E-12 | 2.49E-08 |
| Cluster-33503.83099 | 17.86392 | 118.1429 | -2.7299 | 3.00E-06 | 0.008518 |
| Cluster-33503.82604 | 17.11983 | 106.4645 | -2.6336 | 2.53E-06 | 0.007435 |
| Cluster-33503.99455 | 29.61426 | 168.1658 | -2.5049 | 3.03E-07 | 0.001203 |
| Cluster-33503.98985 | 10642.13 | 58233.13 | -2.452 | 2.11E-12 | 3.73E-08 |
| Cluster-33503.67947 | 46.9285 | 245.385 | -2.388 | 3.29E-07 | 0.001276 |
| Cluster-33503.29591 | 103.148 | 535.5993 | -2.3771 | 7.70E-06 | 0.018372 |
| Cluster-33503.99442 | 159.3264 | 820.5364 | -2.3635 | 1.25E-06 | 0.004152 |
| Cluster-33503.99447 | 226.9377 | 1160.856 | -2.3547 | 5.41E-09 | 3.74E-05 |
| Cluster-33503.54524 | 329.2436 | 1651.53 | -2.3271 | 3.89E-08 | 0.000221 |
| Cluster-33503.74547 | 71.06352 | 343.8941 | -2.2767 | 6.33E-06 | 0.01572 |
| Cluster-33503.50680 | 106.0984 | 468.9649 | -2.1445 | 2.40E-08 | 0.000147 |
| Cluster-33503.127585 | 171.9499 | 758.9011 | -2.1422 | 6.02E-08 | 0.000293 |
| Cluster-33503.142217 | 362.7567 | 1510.353 | -2.0577 | 4.61E-09 | 3.33E-05 |
| Cluster-33503.65963 | 183.4179 | 728.9146 | -1.9904 | 5.16E-08 | 0.000274 |
| Cluster-33503.146366 | 64.33406 | 253.8232 | -1.9797 | 2.11E-05 | 0.042495 |
| Cluster-33503.47433 | 139.3853 | 547.822 | -1.9753 | 5.97E-08 | 0.000293 |
| Cluster-33503.82343 | 95.15411 | 364.9328 | -1.9382 | 7.03E-07 | 0.002523 |
| Cluster-33503.30717 | 139.8437 | 531.9532 | -1.9288 | 1.10E-05 | 0.024208 |
| Cluster-33503.72876 | 1685.051 | 6280.95 | -1.8983 | 1.87E-07 | 0.000761 |
| Cluster-33503.67786 | 388.3833 | 1446.21 | -1.8967 | 6.09E-08 | 0.000293 |
| Cluster-33503.29548 | 300.8745 | 1092.506 | -1.8604 | 5.04E-07 | 0.001906 |
| Cluster-33503.99017 | 840.049 | 3018.019 | -1.8451 | 4.64E-08 | 0.000254 |
| Cluster-33503.99433 | 1030.144 | 3678.842 | -1.8364 | 4.84E-06 | 0.012221 |
| Cluster-33503.100134 | 453.1346 | 1589.09 | -1.8099 | 9.05E-06 | 0.02084 |
| Cluster-33503.27620 | 91.22503 | 317.2852 | -1.7969 | 9.81E-06 | 0.021966 |
| Cluster-33503.98980 | 508.7717 | 1736.197 | -1.7707 | 7.04E-08 | 0.000319 |
| Cluster-33503.99016 | 629.7241 | 2135.446 | -1.7615 | 8.88E-07 | 0.003069 |
| Cluster-33503.93217 | 493.6202 | 1591.578 | -1.6888 | 6.43E-06 | 0.015728 |
| Cluster-33503.122220 | 386.1359 | 1220.254 | -1.6596 | 7.75E-06 | 0.018372 |
| Cluster-33503.101049 | 103.639 | 326.9984 | -1.6574 | 1.72E-05 | 0.035084 |
| Cluster-33503.50332 | 712.5724 | 2184.052 | -1.616 | 1.24E-05 | 0.025979 |
| Cluster-33503.46860 | 1229.407 | 3761.834 | -1.6134 | 3.36E-06 | 0.009362 |
| Cluster-33503.46861 | 216.9503 | 662.6814 | -1.6107 | 2.16E-05 | 0.042999 |
| Cluster-33503.100489 | 129.951 | 380.4445 | -1.5494 | 2.36E-05 | 0.045818 |
| Cluster-33503.93054 | 676.0603 | 1952.662 | -1.5302 | 2.47E-06 | 0.007404 |
| Cluster-33503.75382 | 939.2504 | 317.2232 | 1.5656 | 9.69E-06 | 0.021966 |
| Cluster-33503.35295 | 356.3305 | 83.65786 | 2.092 | 2.22E-05 | 0.043585 |
| Cluster-33503.78577 | 239.3139 | 42.47672 | 2.4952 | 1.21E-05 | 0.025556 |
| Cluster-33503.33273 | 576.0719 | 93.63076 | 2.6223 | 8.61E-06 | 0.020121 |
| Cluster-33503.27714 | 180.362 | 25.62775 | 2.8089 | 1.57E-05 | 0.032501 |
| Cluster-33503.40311 | 7501.224 | 1055.103 | 2.8298 | 1.35E-11 | 1.79E-07 |
| Cluster-33503.35715 | 124.3904 | 16.40437 | 2.9238 | 1.21E-05 | 0.025556 |
| Cluster-33503.76770 | 95.66024 | 10.65527 | 3.1688 | 5.44E-07 | 0.002011 |
| Cluster-33503.84059 | 323.3889 | 18.92224 | 4.0987 | 4.40E-17 | 1.75E-12 |
| Cluster-33503.69414 | 1457.644 | 82.54651 | 4.1441 | 4.51E-22 | 2.39E-17 |
| Cluster-33503.93079 | 64.05922 | 0.682505 | 6.5507 | 8.67E-08 | 0.000383 |
| Cluster-33503.114665 | 111.8005 | 1.026904 | 6.7686 | 7.14E-07 | 0.002523 |
| Cluster-33503.85379 | 99.62864 | 0.701846 | 7.1757 | 1.90E-06 | 0.005821 |
| Cluster-33503.71791 | 710.5572 | 2.449937 | 8.2083 | 1.22E-33 | 1.94E-28 |
| Cluster-33503.74871 | 1137.66 | 0.721187 | 10.697 | 1.26E-22 | 9.98E-18 |
| Cluster-33503.95063 | 186.6346 | 0 | 23.912 | 3.31E-08 | 0.000195 |
| Cluster-33503.96104 | 259.6656 | 0 | 24.361 | 1.82E-08 | 0.000116 |
| Cluster-33503.78173 | 795.2259 | 0 | 25.776 | 2.59E-09 | 2.06E-05 |
| Cluster-33503.103697 | 1325.376 | 0 | 26.565 | 8.36E-10 | 7.00E-06 |
| Cluster-33503.83755 | 1722.539 | 0 | 26.949 | 4.77E-10 | 4.74E-06 |

Supplementary table 5 NCBI accession numbers of *KRT1* sequences from 19 species

|  | Species | | NCBI accession number |
| --- | --- | --- | --- |
|  | *Erinaceus europaeus* | | XM_007535509.2 |
|  | *Odobenus rosmarus* | | XM_004410492.1 |
|  | *Ursus arctos* | | XM_026491053.1 |
| *Ursus maritimus* | | XM_008701257.1 | |
|  | *Ailuropoda melanoleuca* | | XM_011228597.2 |
|  | *Canis lupus* | | XM_025440552.1 |
|  | *Vulpes vulpes* | | XM_025987081.1 |
|  | *Mustela putorius* | | XM_004772755.2 |
|  | *Rousettus aegyptiacus* | | XM_016150993.1 |
|  | *Equus asinus* | | XM_014833554.1 |
|  | *Callorhinus ursinus* | | XM_025861805.1 |
|  | *Condylura cristata* | | XM_004684201.2 |
|  | *Lontra canadensis* | | XM_032871791.1 |
|  | *Phoca vitulina* | | XM_032396498.1 |
|  | *Zalophus californianus* | | >XM_027625808.1 |
|  | *Felis catus* | | XM_006940344.3 |
|  | *Tupaia chinensis* | | XM_006155972.3 |
|  | *Trichechus manatus* | | XM_004390333.2 |
|  | *Callithrix jacchus* | | XM_002748619.3 |
